# Three Species of Axenic Mosquito Larvae Recruit a Shared Core of Bacteria in a Common Garden Experiment

**DOI:** 10.1101/2023.03.23.534051

**Authors:** Josephine Hyde, Doug E Brackney, Blaire Steven

## Abstract

In this study we describe the generation of two new species of axenic mosquito, *Aedes albopictus* and *Aedes triseriatus.* Along with *Aedes aegypti*, axenic larvae of these three species were exposed to an environmental water source to document the assembly of the microbiome in a common garden experiment. Additionally, the larvae were reared either individually or combinatorially with the other species to characterize the effects of co-rearing on the composition of the microbiome. We found that the microbiome of the larvae was composed of a relatively low diversity collection of bacteria from the colonizing water. The abundance of bacteria in the water was a poor predictor for their abundance in the larvae, suggesting the larval microbiome is made up of a subset of relatively rare aquatic bacteria. We found eleven bacterial 16S rRNA gene amplicon sequence variants (ASVs) that were conserved amongst ≥90% of the mosquitoes sampled, including two found in 100% of the larvae, pointing to a conserved core of bacteria capable of colonizing all three species of mosquito. Yet, the abundance of these ASVs varied widely between larvae suggesting individuals harbored largely unique microbiome structures, even if they overlapped in membership. Finally, larvae reared in a tripartite mix of the host species consistently showed a convergence in the structure of their microbiome, indicating that multi-species interactions between hosts potentially lead to shifts in the composition of their respective microbiomes.

**IMPORTANCE:** This study is the first report of the axenic (free of external microbes) rearing of two species of mosquito, *Aedes albopictus* and *Aedes triseriatus*. With our previous report of axenic *Aedes aegypti*, brings the number of axenic species to three. We designed a method to perform a common garden experiment to characterize the bacteria the three species of axenic larvae assemble from their surroundings. Furthermore, species could be reared in isolation or in multi-species combinations to assess how host species interactions influence the composition of the microbiome. We found all three species recruited a common core of bacteria from their rearing water, with a large contingent of rare and sporadically detected bacteria. Finally, we also show that co-rearing of mosquito larvae leads to a coalescence in the composition of their microbiome, indicating that host species interactions potentially influence the composition of the microbiome.

## INTRODUCTION

Many organisms harbor a community of associated microbes collectively referred to as the microbiome. Understanding how the microorganisms associated with the host are recruited out of the environment remains a significant challenge for host/microbiome studies (1–3). Mosquitoes are thought to largely acquire the microbes that make up their microbiome from the aquatic environment in which they are reared (4–6). Furthermore, there is some evidence that different mosquito species assemble disparate microbiomes (7, 8), even if they are reared in the same environment (9). Yet, these data are often confounded with the date and location of mosquito collection, or the age and health of the mosquito, all of which have been shown to influence the composition of the microbiome (10–13). Thus, these observations support that the assembly of the microbiome in the mosquito is complex and dependent on a variety of host and environmental factors.

A substantial hurdle to developing mechanistic models for microbiome assembly is the lack of systems in which the microbiome can be manipulated. In 2018, we reported the generation of an axenic *Aedes aegypti* mosquito model, demonstrating that a living microbiome is not necessary for mosquito development (14). Importantly, the axenic model is a blank template that can be used to study the bacteria that colonize the host, opening an avenue for manipulative studies (15). This includes methods to introduce environmental bacteria, in a process we refer to as ‘rewilding’ the microbiome (16). Here we describe the axenic rearing of two additional species of mosquito, *Aedes albopictus* (Asian tiger mosquito) and *Aedes (Ochlerotatus) triseriatus* (Eastern tree hole mosquito).

Of the three mosquito species, *Ae. albopictus* has the largest geographic range, spanning most of the temperate and tropical globe, whereas *Ae. aegypti* is primarily found in tropical and subtropical regions (17). *Ae. triseriatus* is the most geographically restricted, being limited to eastern and central North America (18). Additionally, each of these mosquito species are known vectors for numerous arthropod-borne viruses, including dengue virus and Zika virus for *Ae. aegypti* and *Ae. albopictus*, and LaCrosse virus is vectored by *Ae. triseriatus*. While *Ae. aegypti* is highly anthropophilic, breeding in man-made containers in and around households, it can also breed in natural habitats outside the home (i.e. tree holes (19, 20)). Both *Ae. albopictus* and *Ae. triseriatus* will similarly breed in both natural and man-made habitats but they are less associated with humans and prefer vegetated urban, suburban, and rural landscapes and forests (21, 22). Due to their geographic and ecologic overlap it is possible to have all three species co-exist in certain locales. In fact, larvae from all three species were found cohabiting discarded tires in Texas (23). Thus, these mosquito species are a model system to interrogate how interacting species differ in the structure of their microbiomes.

To address the assembly of the microbiome among these different species of mosquito, axenic larvae were exposed to an identical source of colonizing water, and we characterized the bacteria that were recruited to the microbiome in a common garden experiment. In this manner, we could document how age synchronized larvae, reared in identical environmental conditions, and exposed to the same source of colonizing bacteria assembled their microbiome. Furthermore, by rearing the mosquitoes either individually or in a common garden we could evaluate how co-rearing of mosquito larvae may influence the composition of their microbiome. We hypothesized that mosquitoes reared individually would harbor a more distinct and variable microbiome in comparison to their co-reared counterparts as they were essentially stranded with the bacteria they collected from the colonizing water. In contrast, co-reared individuals had the potential enrich and transfer bacteria between larvae. In this vein, we also predicted that if the different mosquito species did assemble different microbiomes, some of those differences might be ameliorated when they were co-reared, as bacteria able to colonize multiple mosquito species would be enriched and shared amongst species. Thus, by directly controlling for a suite of host and environmental factors we could identify the bacteria larvae were recruiting from their surroundings as well as interrogate the influence of a shared rearing environment on the composition of the microbiome.

## RESULTS

### Growth of axenic mosquitoes

Here we report the axenic rearing of two species of mosquito from larvae to adult, *Ae. albopictus* and *Ae. triseriatus,* using the protocol we previously described for *Ae. aegypti* (14). Both species were able to develop to adulthood when raised axenically, however they both experienced developmental delays compared to larvae raised conventionally, 5.3 days on average for *Ae. albopictus* and 7.8 days for *Ae. triseriatus* (Figure 1). This matches our previous report for *Ae. aegypti,* suggesting that axenic rearing is associated with developmental delays but underpinning that live bacteria are not required for mosquito development.

**Figure 1.**
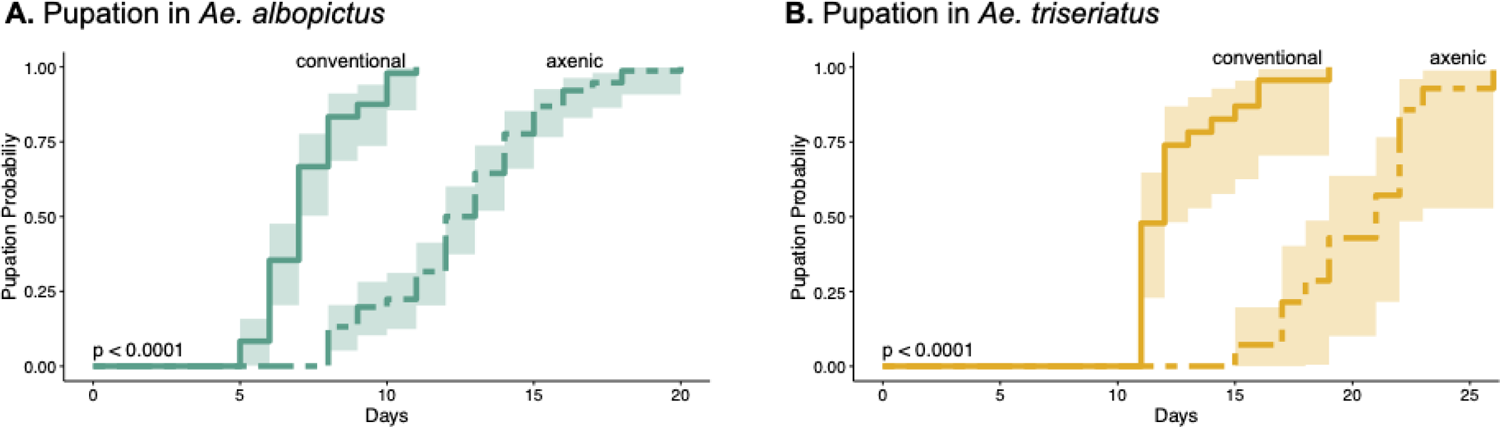
Rearing of axenic *Ae. albopictus* and *Ae*. *triseriatus*. The probability of pupation for each species per day is shown for conventionally reared and axenic larvae. The shaded boxes denote the 95% confidence intervals. Axenic larvae were shown to have a significant delay in time to pupation via a cox regression test. The p-value of the test statistic is indicated in the lower left corner.

### Experimental design

To characterize the bacterial taxa that larvae assembled from their aquatic environment, axenic larvae were exposed to an environmental water source simulating a mosquito breeding site. The axenic larvae were exposed to the water for 2.5 hours and then transferred individually into four different rearing conditions (Figure 2). **Individual**, single larvae were transferred to a well of a 6-well plate so that they could not exchange microbes between any other mosquitoes. **Self,** individual larvae were transferred to a sterile rearing cup allowing for the exchange of microbes between larvae of the same species (Figure 2). **Paired,** larvae were reared in each pairwise combination allowing for microbial exchange between species. **Tripartite**, larvae were reared in a mixture of all three species. Larvae were allowed to develop for four days, at which point they were harvested, and their microbiome was characterized *via* 16S rRNA gene sequencing.

**Figure 2.**
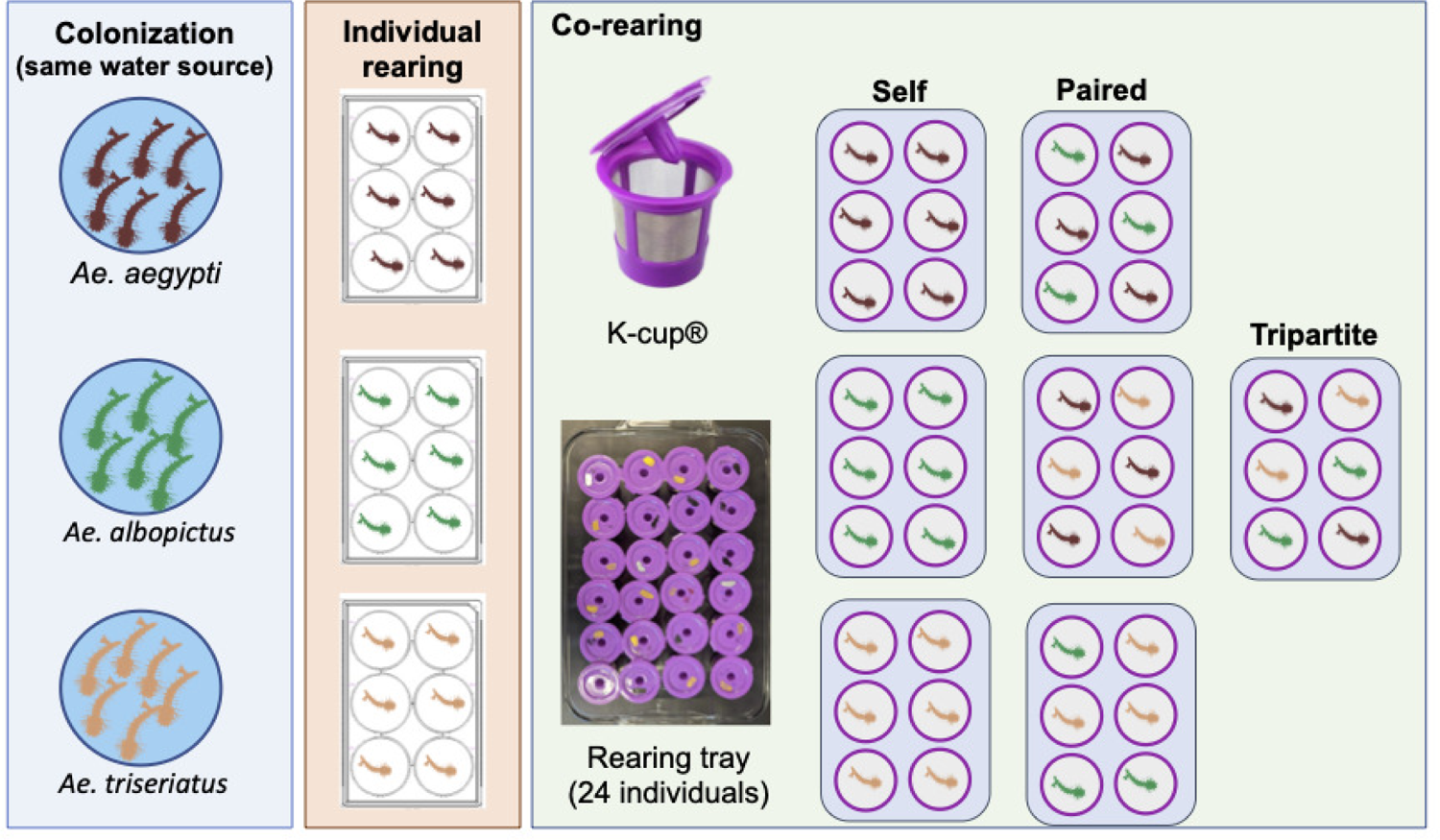
Schematic diagram of the experimental setup. Axenic larvae of the three species were hatched in aliquots of an environmental water source in petri dishes. The larvae were left in the water for 2.5 hours to allow for bacterial colonization. After colonization, single larvae were transferred to wells of a 6-well plate for **individual** rearing. These larvae were unable to exchange microbes with any other individuals. For the co-rearing experiments, single larvae were placed in an autoclaved K-cup. We hypothesized that the filters would allow for the exchange of microbes between individuals. Co-rearing conditions consisted of **self**, larvae reared with other members of the same species; **paired**, each combinatorial group of two species mixtures; and **tripartite,** a mixture of all three species. In each mixed condition larvae were randomly distributed. The co-rearing experiments had 24-individuals per tray, but only six are shown in the figure for ease of viewing.

### Mosquitoes recruit a subset of rare bacteria out of the colonizing water

Our first objective in the common garden experiment was to determine which bacteria the mosquitoes were recruiting out of the colonizing water. Non-metric Multidimensional Scaling (NMDS) ordination showed a clear clustering of the 16S rRNA gene datasets from the mosquitoes separately from water samples collected at the time of mosquito colonization (Fig. 3A). The clustering was statistically supported with a Permutational Multivariate Analysis of Variance (PERMANOVA) p-value of <0.001. These observations indicate the mosquito assembled a microbiome that was distinct from the initial water source. In addition, Shannon’s diversity for the sequence datasets revealed a significantly higher alpha diversity for the water samples than sequences recovered from the larvae, further supporting a difference in the composition of the microbial populations inhabiting the mosquitoes, pointing to a less diverse community in the larvae (Fig, 3B). Yet, there was no significant differences in alpha diversity among 16S rRNA gene datasets from the mosquito species, suggesting diversity of the microbiome was similar across the host species (Fig. 3B).

**Figure 3.**
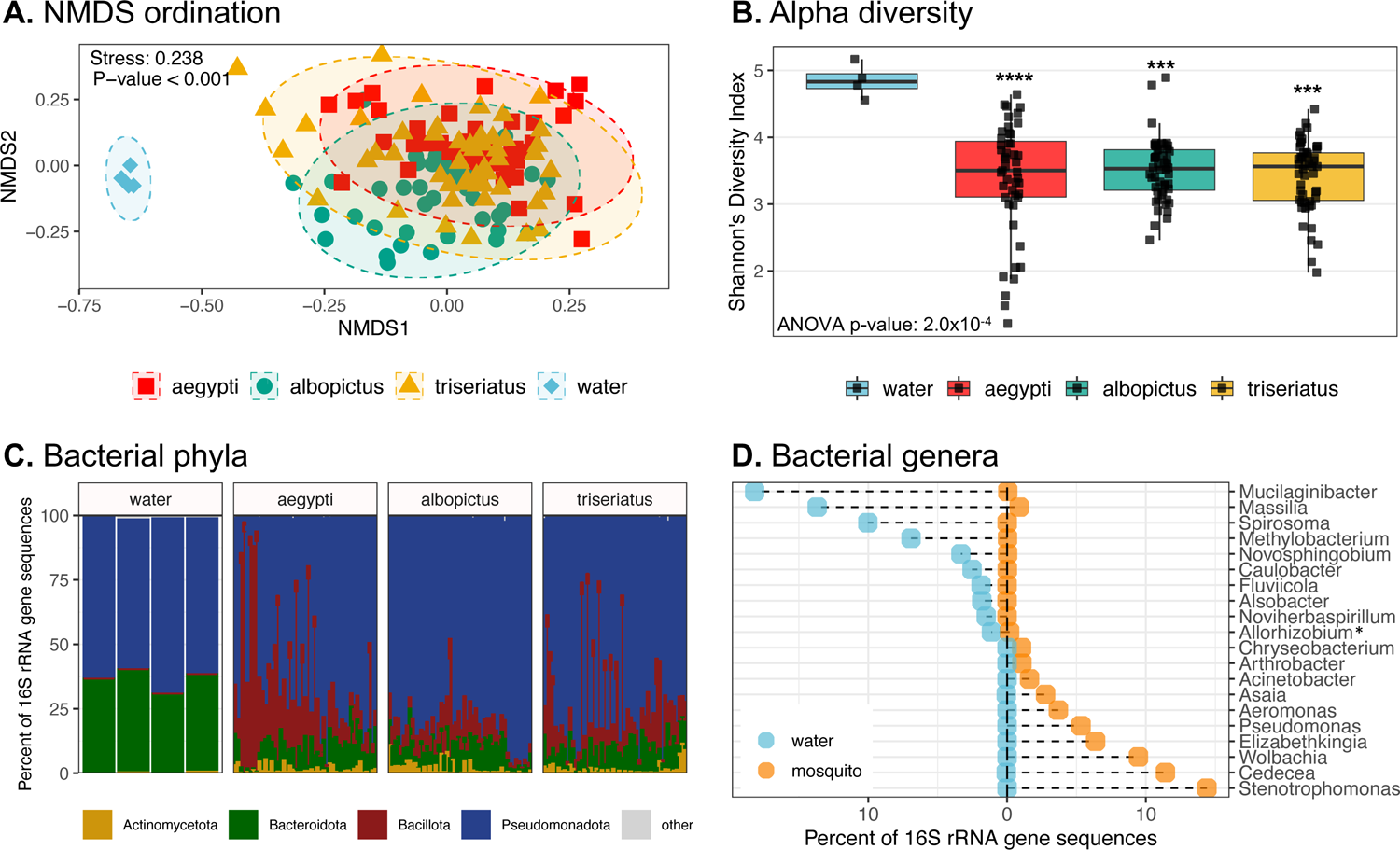
Recruitment of bacteria from colonizing water. **A)**. NMDS ordination. Inter-sample distances were calculated with the Bray-Curtis metric. Ellipses denote the 95% confidence level for the distribution of each group. The stress value for the ordination is indicated along with the p-value from a PERMANOVA test. **B)**. Alpha diversity. Shannon’s diversity index was calculated for each sample. The ANOVA p-value from comparisons of means is indicated and the asterisks indicate the p-values of a post-hoc t-test for comparisons between datasets from the three mosquito species compared to the water samples. P-value key **** = p <0.0001, *** = p<0.001. Comparisons between mosquito species were not significant. **C)**. Relative abundance of phylum level bins in the datasets. Only the four most abundant phyla are shown with the remainder assigned to the catagory “other”. **D)**. Conservation of bacterial genera between water and mosquitoes. The 20 most abundant genera in the dataset are displayed. Each point represents sequences assigned to a bacterial genus and the mean abundance of that genus in either water samples or mosquito larvae. Only sequences classified to the rank of genus are shown. *The genus Allorhizobium is shortened from the SILVA label Allorhizobium-Neorhizobium-Pararhizobium-Rhizobium.

To investigate the taxonomic composition of the bacterial communities, 16S rRNA gene sequences were classified to the phylum level (Fig. 3C). Observationally, the proportion of the phylum Bacteroidota was higher in the water samples (mean 37%) compared to mosquitoes (mean 8%). Conversely, the phylum Bacillota was elevated in the mosquitoes (mean 18%) while only accounting for <1% of sequences in the colonizing water. Similarly, at the rank of genus, the mean abundance of the numerically dominant taxa in both the water and mosquitoes were plotted (Fig. 3D). Overall, the most abundant genera in the water were not particularly abundant in the mosquitoes and *vice versa*. For instance, the most abundant genera in the water were *Mucilangibacter*, *Massilia*, and *Spirosoma* accounting for 18%, 13%, and 10% of sequences, respectively. Yet, in the mosquitoes these genera made up less than 1% of 16S rRNA gene sequences collectively. Similarly, the genera *Stenotrophomonas* and *Cedecea*, which were most abundant in the mosquitoes (14% and 11% of sequences), were only detected in the water with abundances of 0.01 and 0.05%, respectively. Thus, these data support that mosquitoes are not simply recruiting the most abundant bacteria from the local species pool, rather they select and enrich a lower diversity set of specific bacterial taxa that are presumably adapted to inhabiting the larvae.

### A core microbiome shared between mosquito species

We next wanted to determine if the microbes recruited to the mosquito microbiome were shared amongst the three mosquito species. The one hundred most abundant 16S rRNA gene ASVs were plotted in a presence-absence heatmap to investigate their conservation amongst individual mosquitoes (Fig. 4A). Most ASVs were only sporadically detected in the microbiomes, being recovered from some individuals but not others. There did not appear to be a large effect of host species on the presence of ASVs as most of the ASVs were detected in individuals of three species.

**Figure 4.**
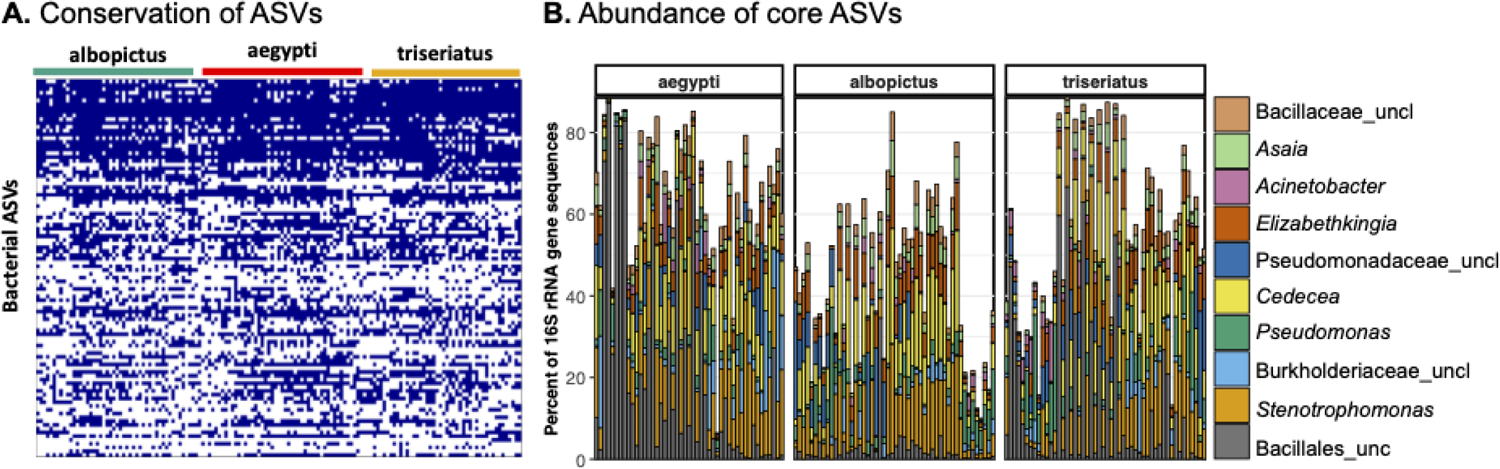
Core bacterial taxa in the mosquito microbiome. **A)**. Presence-absence heatmap of bacterial ASVs among mosquito species. Each column represents an individual mosquito. Presence of the ASV is indicated in blue. The 100 most abundant ASVs in the sequence dataset are displayed. **B).** Relative abundance of the 11 conserved ASVs in individual mosquitoes. Bars are coloured by the deepest classification for the ASV (Table 2). Because two ASV’s were classified the genus *Stentrophomonas* bars represent the sum of both ASVs.

Notably, we did detect two ASVs, classified to the genera *Stenotrophomonas* and *Elizabethkingia*, that were identified in all mosquitoes assayed (Table 1). When the requirements for conserved ASVs was relaxed to those taxa that were conserved amongst at least 90% of mosquitoes, the number of ASVs increased to eleven, and included genera such as *Acinetobacter*, *Asaia*, *Cedecea*, *Pseudomonas,* and another ASV within the genus *Stenotrophomonas*. Three of the ASVs could not be classified to the genus level, being classified to the order *Bacillus*, another to the family *Bacillaceae*, and the final ASV to the family *Burkholderiaceae* (Table 1). Thus, these taxa represent a ‘core’ microbiome representing bacterial taxa that were present in a large majority of the mosquitoes.

**Table 1.**
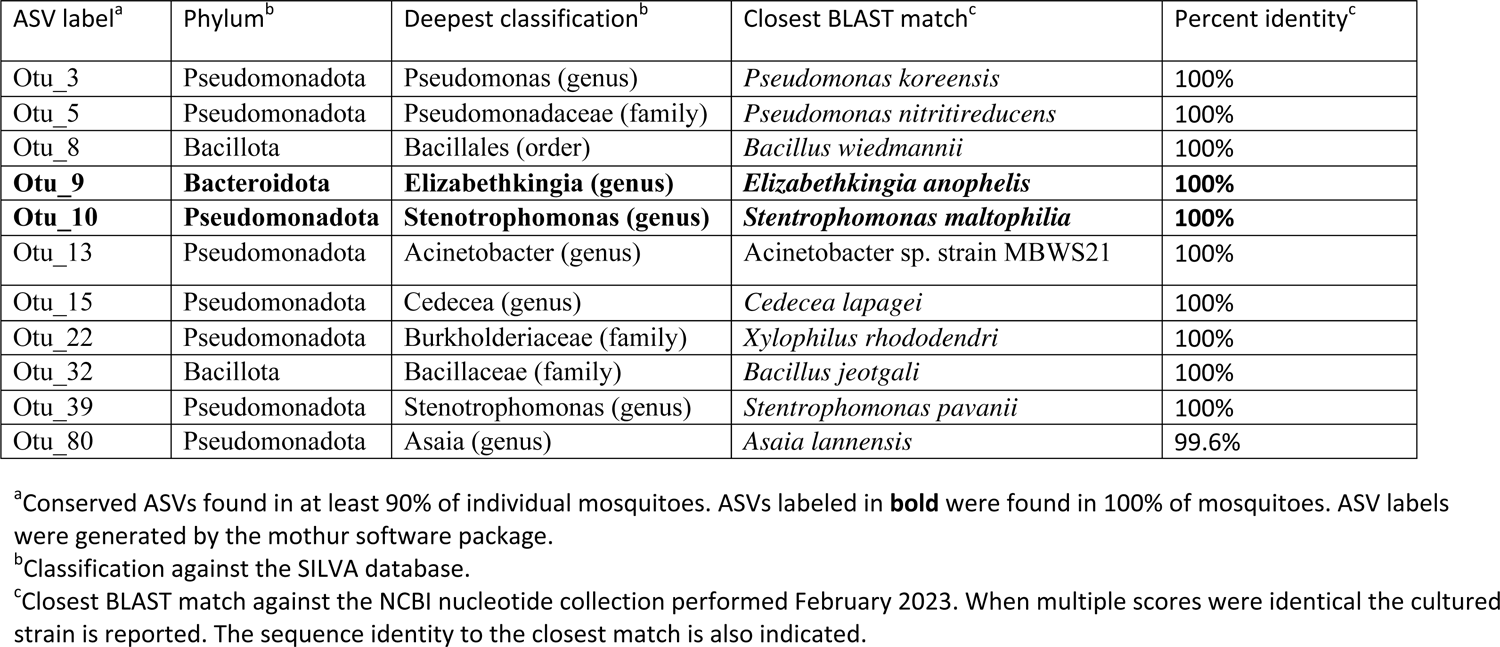
Identification of conserved ASVs

We next examined the proportional abundance of these core ASVs in the microbiomes to test if they were conserved in both presence and relative abundance. The relative abundance of the ASVs varied widely amongst individual mosquitoes (Fig. 4B). The combined abundance of these 11 ASVs ranged from >90% of the recovered sequences to <10%. Similarly, the individual ASVs varied in relative abundance between species. As a single exemplar, the ASVs identified as *Stenotrophomonas* had a relative abundance range of ∼1% to >30%, despite being present in every mosquito assayed. In this regard, at the level of abundance, individual mosquitoes largely possess a unique microbiome structure, even if the members of the community are overlapping.

Having shown that the mosquitoes share a common core of bacterial ASVs does not answer the question as to whether the composition of the microbiome differs amongst host species. For example, the microbiome may be differentiated by rare taxa or at a higher taxonomic rank. To address this, we identified significantly differentially abundant ASVs amongst the host species. Only two ASVs were identified as differentially abundant, and both were significantly enriched in *Ae. albopictus* (Fig. 5A; supplemental Table 1). Both ASVs were identified as belonging to the genus *Wolbachia*. To further investigate these observations, the relative abundance of sequences classified to the genus *Wolbachia* are plotted in Figure 5B. *Wolbachia*-related sequences were identified in all of the *Ae. albopictus* larvae, with relative abundances ranging from 0.3% to 70% of recovered 16S rRNA sequences (mean 24%). *Wolbachia* sequences were detected in the majority of *Ae. aegypti* (96%) and *Ae. triseriatus* (92%) larvae, but the relative abundance tended to be much lower than for *Ae. albopictus.* The maximal abundance of *Wolbachia*-related sequences was 14% in *Ae. aegypti* and 11% in *Ae. triseriatus* larvae. Thus, these data support that the colony *Ae. albopictus* mosquitoes are largely subject to *Wolbachia* infections. In contrast, the status of *Wolbachia* in *Ae. aegypti* and *Ae. triseriatus* is less certain. While *Wolbachia* was detected in the majority of individuals, it is not established that these represent true infections. Importantly, the presence of *Wolbachia* may be important to the biology of these organisms, as *Wolbachia* are a group of intracellular endosymbionts possessed by a wide range of insects, with potential roles in the reproduction and fecundity of the host organism (e.g. 16– 18). Here we show all three species are potentially susceptible to *Wolbachia* infection, but the infection rates and burdens were significantly higher for *Ae. albopictus*. Importantly, *Wolbachia* sequences were never detected in the water, supporting these are vertically transmitted intracellular bacteria and were not likely to have been obtained from the source of colonizing water.

**Figure 5.**
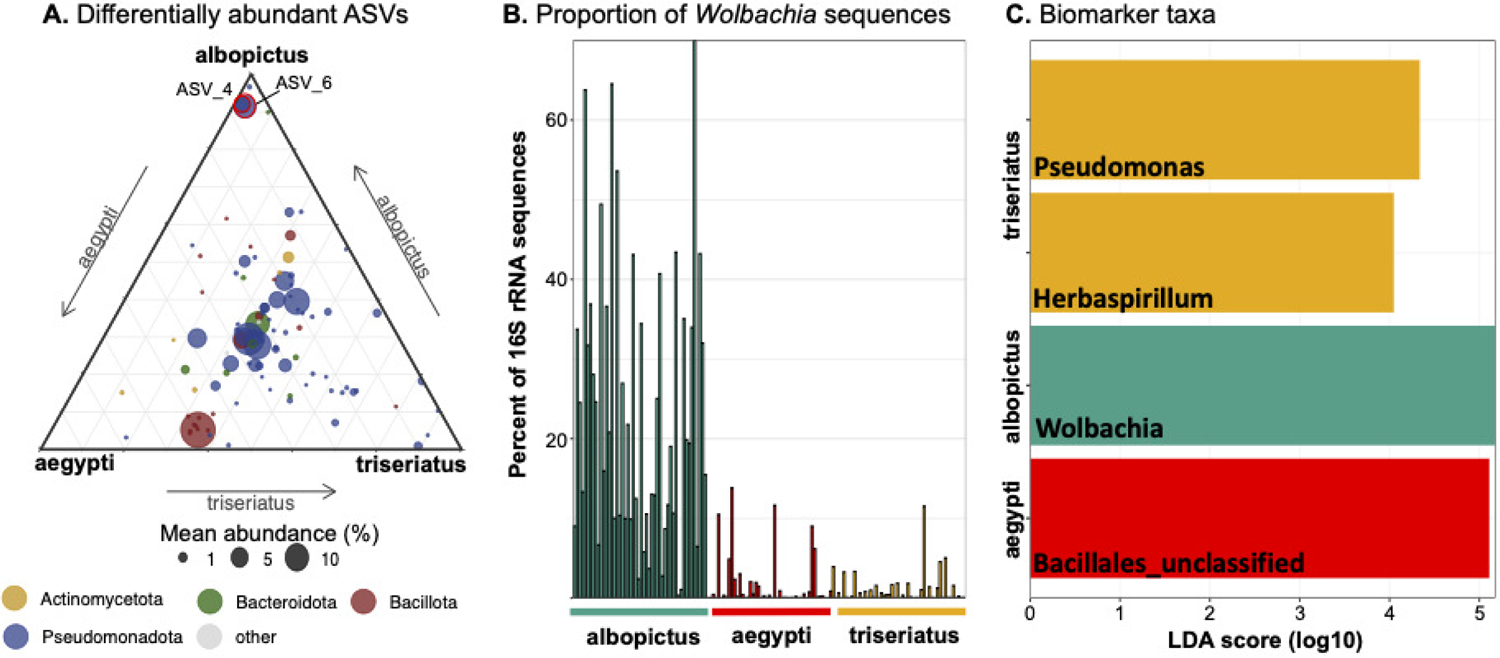
Differentially abundant bacterial taxa among mosquito species. **A)**. Ternary diagram showing the mean relative abundance of ASVs in each mosquito species. ASVs with a significant difference in abundance are shown in red. The data for the significance test are shown in supplemental Table 1. Both ASVs were classified to the genus *Wolbachia* and were enriched in *Ae. albopictus*. **B)**. Proportion of *Wolbachia* sequences in mosquitoes. Each bar represents an individual larva. **C**). LDA effect size (LEfSe) analysis at the genus level among mosquito species. Only categories meeting a log linear discriminant analysis (LDA) significant threshold >4 are shown. Bars are coloured by the mosquito species in which the genera were enriched.

Finally, a Linear discriminant analysis Effect Size (LEfSe) analysis was performed to identify any potential biomarker taxa that may distinguish the microbiome status amongst the host mosquito species. Unsurprisingly, this analysis identified an enrichment of the genus *Wolbachia* as a potential biomarker for *Ae. albopictus*, however it also identified elevated abundances of the taxa *Pseudomonas* and *Herbaspirillum* as potential biomarker strains for *Ae. triseriatus* and bacteria in the order *Bacillales* as potentially diagnostic of the *Ae. aegypti* microbiome.

Taken together, these analyses point to only few bacterial taxa potentially differentiating the microbiomes of three species of mosquito. Notably, the most diagnostic bacteria that differentiated the microbiomes was an intracellular endosymbiont that was unlikely to have arisen from the shared colonization water. Instead, these three species of mosquito recruited a small group of core taxa, along with a much larger contingent of bacteria that were only sporadically detected in the microbiome, with no substantial pattern of conservation or abundance among any particular host species.

### Influence of co-rearing on the microbiome

The final objective of this study was to document how co-rearing of mosquitoes would influence the composition of the microbiome. NMDS clustering of the datasets consistently retuned a PERMANOVA p-value <0.001, indicating that co-rearing resulted in a different microbiome composition across the species (Figure 6). Extracting the pairwise dissimilarity values showed that co-rearing of species almost always decreased the dissimilarity scores in comparison to the individually reared mosquitoes, with the sole exception of *Ae. albopictus* raised in the presence of *Ae. aegypti* (Figure 6). The effect was particularly apparent in the tripartite rearing, in which community dissimilarity was consistently lower than for individual or self-reared mosquitoes. Thus, co-rearing of mosquito species appears to be a homogenizing influence on the mosquito microbiome, producing individuals with more similar community composition. In other words, individuals of the same mosquito species had a more similar microbiomes when they were reared in a tripartite host species mix compared to when they were raised individually, or even with other members of their own species.

**Figure 6.**
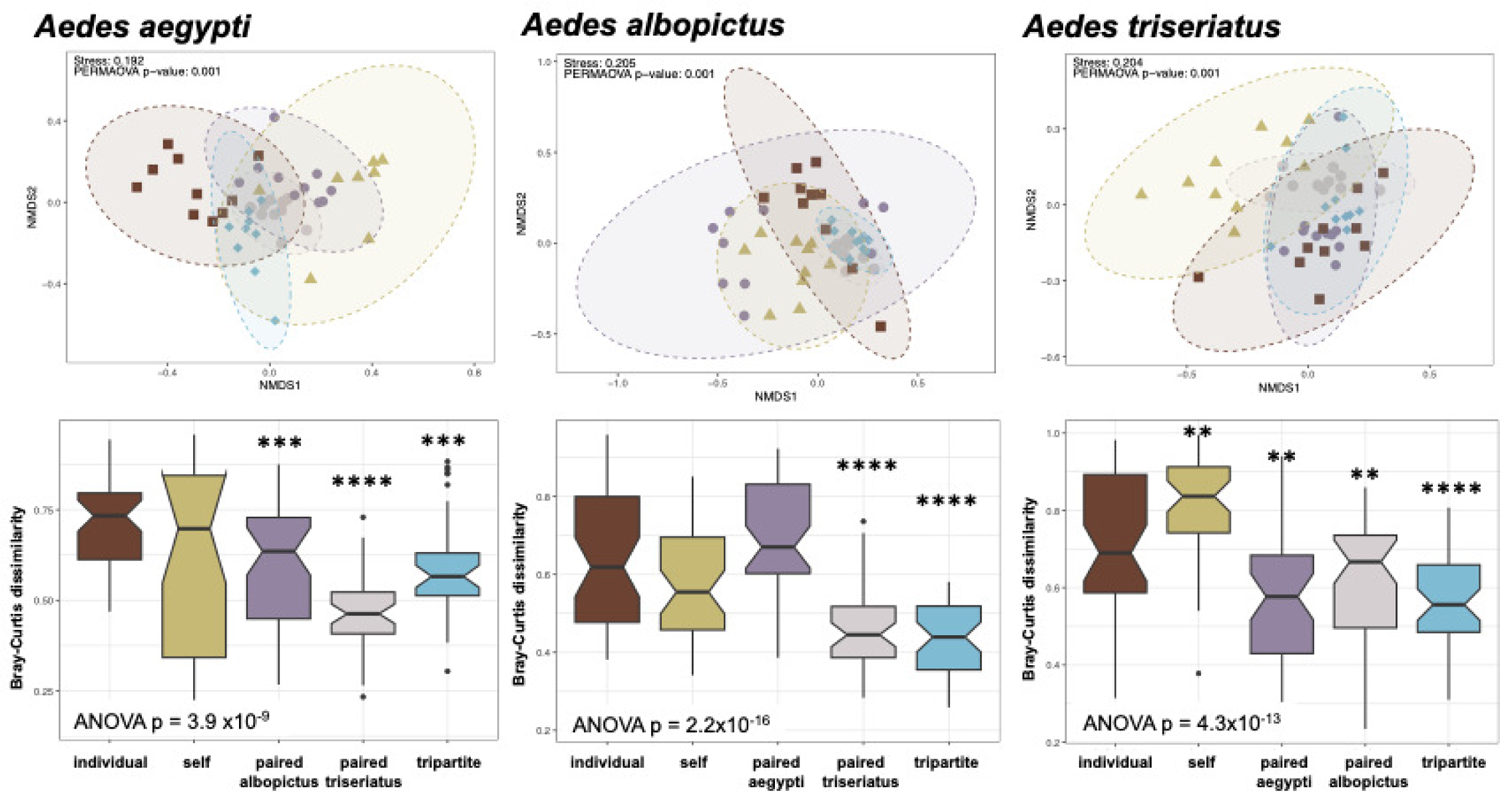
Co-rearing of mosquito larvae. Each NMDS plot represents individual larvae raised in the different rearing conditions. *Individual*, reared in isolation; *self,* reared with members of the same species; *paired*, raised with another species; *tripartite*, raised with all three species. Inter-sample distances were calculated with the Bray-Curtis metric. Ellipses denote the 95% confidence level for the distribution of each group. The stress value for each ordination is indicated along with the p-value from a PERMANOVA test. The violin plots below are the pairwise dissimilarity measures (Bray-Curtis dissimilarity) between individuals in each condition. The ANOVA p-value from comparisons of means is indicated and the asterisks indicate the p-values of a post-hoc t-test for comparisons between datasets from the co-rearing conditions with the samples from individual rearing conditions set as a reference level. P-value key **** = p <0.0001, *** = p<0.001, **=p<0.01.

We additionally tested to determine if any bacteria taxa were differentially abundant due to co-rearing, and found a set of 6, 9, and 10 genera in *Ae. aegypti*, *Ae. albopictus*, and Ae. *triseriatus*, respectively (supplemental Table 2-4). Yet, none of these genera were identified as differentially abundant across all three host species. This suggests that the convergence in microbiome structure under co-rearing is not explained by the enrichment of depletion of a particular bacterial taxon, rather appears to be largely a host specific response.

## DISCUSSION

In this study, we employed a common garden experiment to document how three species of microbially naive mosquitoes assemble their microbiome out of a common pool of colonizing bacteria. Relative to the initial water samples, the mosquitoes were enriched for the phyla Bacillota and Actinomycetota (Figure 3C), more specifically genera such as *Stentrophomonas, Cedecea,* and *Elizabethkingia* showed an asymmetrical distribution, being more abundant in the mosquitoes than in the colonizing water (Figure 3D). These data suggest that mosquitoes are selectively enriching a subset of bacteria out of the larger and more diverse species pool that is present in the colonizing water (Figure 3B). It is well accepted that hosts appear to ‘choose’ appropriate microbial partners, but understanding the characteristics of the bacteria that make for successful colonizers remains elusive. Certain bacterial traits such as a rapid growth rate, provisioning the host with essential nutrients, or increasing the host’s phenotypic plasticity by allowing it to diversify its diet or tolerate environmental change have been identified as characteristics that may favor host colonization (27–29). Additionally, the capacity to synthesize bioactive compounds that provide the host with defensive measures against predators, parasites and pathogenic microorganisms seems to enhance host colonization (27, 30). For example, the production of antibiotics by microbiome members may benefit the host by excluding pathogens, but may also benefit the microbe itself by giving it a competitive advantage in the microbiome (31). We have previously shown that antibiotic resistant bacteria are common in both field-caught mosquitoes and colony reared *Ae. aegypti* larvae and adults, suggesting that antibiotic production and defence may be common in the mosquito microbiome (32). Connectedly, bacteria in the genera *Elizabethkingia* and *Stentrophomonas*, which were both identified as core bacterial taxa (Table 1), were also identified as antibiotic resistant bacteria in our previous screen (32). Thus, these observations highlight the power of these common garden experiments to identify the bacteria that excel at colonizing mosquitoes and to begin to assess the traits that predict their colonization success.

We were also able to assess the microbiomes assembled by three species of mosquito. Many studies have been conducted to address host specificity in microbiome assembly. Examples abound of both strong influences of the host on microbiome structure, including in mosquitoes (33–35) and counter examples of host species only playing a limited role in microbiome assembly (36–38). Yet, when the relative influence of the environment versus host have been compared, the effects of the environment usually dominate (39–41). By controlling the environmental variables in a common garden experiment, we found that three species of axenic hosts gather a shared core of microbial taxa, suggesting that there was not a strong host filtering between bacterial taxa. In this respect, when the externalities of environmental variables and host age are removed, these three species of mosquitoes largely assemble comparable microbiomes. The finding of similar microbiomes between species in this experiment may be at least partially due to an experimental artifact. Mosquitoes have a potential avenue for the vertical transmission of bacteria from mother to offspring in the process of “egg smearing”, where bacteria colonize the mosquito ovaries and are transmitted to larvae on the surface of eggs (42). As our method to produce axenic larvae involves the surface sterilization of eggs, removing any adhering bacteria, this route of transmission is blocked. In *Drosophila* the microbiome of axenic flies was dominated by bacteria most abundant on the eggs, demonstrating effective vertical transmission (43). Thus, this may be a mechanism by which different species of mosquito maintain different microbiomes, in that microbes are vertically transmitted, rather than acquired from the environment. Future studies including alternative methods for the acquisition of the microbiome, or multi-generational common garden experiments may shed light on the relative contribution of vertical transmission on the establishment of the microbiome.

To the extent that we did see any differences in microbiome structure between the host species, it was primarily driven by the high infection rates of *Wolbachia* in the *Ae. albopictus* mosquitoes (Figures 5B,C). As mentioned previously, *Wolbachia* is an endosymbiont that is vertically transmitted and thus unlikely to have arisen from the colonizing water. However, there is no consensus on whether *Wolbachia* infection influences the assembly of the mosquito microbiome, with studies reporting both significant influences (44, 45) or no impact (46, 47). This is potentially important as the structure of the microbiome has been reported to influence the vertical transmission and biology of *Wolbachia* (48). Thus, uncovering potential *Wolbachia-*microbiome interactions is an acute research need. In this study we show that *Ae. albopictus* which harbors a significantly higher *Wolbachia* infection rate and burden than either *Ae. albopictus* or *Ae. triseriatus* recruits a largely similar microbiome, indicating that *Wolbachia* infection did not seem to play a significant role in microbiome assembly. This demonstrated the utility of having three species of axenic mosquitoes, as it opens a window to mechanistically study host microbiome interactions in *Wolbachia* permissive and resistant mosquitoes. Additionally, it is possible to cure organisms of *Wolbachia* (49), opening the potential of a truly axenic strain, free from intracellular bacteria, and further increasing the capacity to employ these axenic strains to study mosquito-*Wolbachia*-microbiome interactions.

Rather than host specific microbiomes, we largely found a core set of bacteria conserved amongst mosquitoes. Importantly, this is not the first time a ‘core microbiome’ has been described for mosquitoes, suggesting there exist many microbes that can be classified as mosquito generalists (50–52). The core taxa identified in this study included members of the genera *Stenotrophomonas* and *Elizabethkingia* which were conserved amongst all larvae surveyed (Table 1). Both taxa have been previously reported to colonize mosquitoes (53–55), in fact, the type species for the genus *Elizabethkingia* was initially isolated from the *Anopheles gambiae* mosquito (56). Other core taxa, found in the majority of mosquitoes included*, Acinetobacter*, *Asaia*, *Cedecea*, *Pseudomonas,* also known to form associations with mosquitoes from a variety of species (42, 57–59). However, it is important to note that much larger than the number of core taxa conserved between mosquitoes was the number of taxa that were rarely or only sporadically detected in the microbiome (Figure 4A). Each mosquito was associated with tens to hundreds of ASVs that fell into this so called ‘rare biosphere’ (12, 60). Increasingly, rare species are found to play an outsized role in microbial ecosystems, driving nutrient cycling and potentially contributing to the host’s phenotypic plasticity (61, 62). Thus, the bacteria that were divergent among the mosquitoes may be as important, if not more important, than the conserved ASVs.

The final mechanism that we investigated as a factor influencing the assembly of the microbiome was the transmission of microbes between individuals and species. For instance, people sharing a home have been shown to share common bacteria with each other, along with their pets (63, 64). Thus, a common household can be a strong determinant in microbiome structure. Here we show that when three species of mosquito are co-reared in the same common garden, they experience a coalescence in their microbiome structure in comparison to mosquitoes reared in isolation or with individuals of the same species (Figure 6). This matches observations in zebra fish, where dispersal of microorganisms among hosts was often larger than the effects of individual host factors on the composition of the microbiome (65). The authors of the zebra fish study posit that microbiome homogenization occurs because the fitness landscape changes for the microbial populations, selecting bacteria with traits for motility, transmission, and colonizing organisms with different genetic backgrounds (65). While we did not identify any specific bacterial taxa that accounted for the homogenization of the mosquito microbiome, we posit that this experimental setup could be expanded to address these questions in a more mechanistic manner. More importantly, these data highlight that the organisms that share a habitat are potentially all part of the same microbial pool, exchanging microbes and co-influencing the composition of their respective microbiomes. In this respect, no microbiome is an island and can be influenced by a variety of environmental factors, including the other host organisms it shares a habitat with. These observations support the importance axenic hosts and experimental designs scaled to disentangle the myriad of host and environmental factors that contribute to the composition of the microbiome.

## MATERIALS AND METHODS

### Developing axenic mosquitoes

Insectary reared mosquitoes of *Ae. albopictus* (colony established 2014) and *Ae. triseriatus* (colony established 1992) were treated as previously described to produce axenic *Ae. aegypti* (14, 66). Briefly, eggs were collected from colony mosquitoes, and in a sterile biosafety hood were serially rinsed for 12 minutes in 70% ethanol, followed by a five-minute wash in a 3% bleach and 0.2% ROCCAL-D (Pfizer) solution, and then again for five minutes in fresh 70% ethanol. The sterilised eggs were then rinsed three times in autoclaved DI water and placed in a Petri dish filled with phosphate-buffered saline (PBS). Eggs were hatched in a vacuum oven (Precision Scientific, Model 29) at 25Hz for 15 minutes at room temperature, producing age synchronized larvae.

Single larvae were transferred from the petri dish to individual wells of six well tissue culture plates containing 5 ml of sterilised DI water and a 0.6 g plug of liver yeast agar (66). Larvae from each mosquito species were split into two groups; axenic receiving no bacterial inoculation and conventional, which received a 10 µl aliquot of a homogenized colony larva from the same species as a source of colonizing bacteria. The larvae were then assessed for their success in morphogenesis and time to pupation. Sterility of the axenic group was verified by culturing viable bacteria and 16S rRNA gene PCR as previously described (14).

### Common garden experiment

To obtain relevant environmental bacteria for colonizing the larvae, we generated mosquito breeding water by filling two 10L polyethylene storage containers with tap water and leaf litter and covered the top with netting to limit inputs. The water was left to stagnate for 2 weeks. Immediately preceding the collection of colony mosquito eggs, the seasoned water was taken from the containers and filtered through autoclaved cheesecloth to remove any large particulates, and one litre was stored in a sterile pyrex media storage bottle. These steps all occurred outside the lab to prevent the introduction of any laboratory bacteria into the experiment.

Eggs from the three different mosquito species *Ae. aegypti*, *Ae. albopictus*, and *Ae. triseriatus* were surface sterilised using the method described above. Newly hatched larvae of each species were transferred to a sterile petri dish containing ∼20 ml of the mosquito breeding water and the larvae were incubated for two and a half hours to acquire colonizing bacteria. Following colonization individual larvae were collected from the petri dishes with a 100 µl pipette set to 10 µl and transferred either to a single well of a six-well culture plate for individual rearing or to a K-Cup® filter for the co-rearing experiments. The K-Cups allowed for exchange of bacteria between mosquitoes, whereas the single larvae in six well plates were reared in isolation. The larvae were reared in the K-Cups to facilitate identifying larval species in the mixed conditions and to keep the larger larvae predating the smaller larvae (9). A schematic diagram of the experimental design is shown in Figure 2.

After the transfer of larvae, the remaining colonization water was vacuumed filtered through a 0.8 µm isopore membrane filter. Two papers were used and stored in a sterile petri dish at −80 °C for DNA extraction and bacterial 16S rRNA gene sequencing. Each filter was split in half prior to DNA extraction, resulting in 4 water samples to characterize the bacterial composition of the colonization water. DNA was extracted directly from the filter by placing the filter in a DNeasy PowerSoil Kit (Qiagen) bead beating tube and following the manufactures directions.

Individual larvae reared in six well culture plates received 5 ml of autoclaved DI water and 100 µL of a 2% liver powder:yeast extract (3:2 ratio) solution. Each experiment consisted of 4 six-well plates (24 individually reared larvae). In the co-rearing experiments, 24 K-Cups were placed in an autoclaved Pyrex tray containing 1 L of autoclaved DI water with 20 mL of 2% liver powder:yeast extract, and covered. All treatment groups were maintained in an environmental chamber at 28 °C 70% relative humidity with a 16:8 light:dark photoperiod *16S rRNA gene amplicon sequencing and bioinformatics* After four days of rearing, ten individual mosquitoes, per species, per treatment group were harvested. The larvae were serially washed in sterile H_2_0, and then frozen at −80 °C until DNA extraction. The larval microbiome and the initial water community on membrane filters, were subjected to DNA extraction using the DNeasy PowerSoil Kit (Qiagen). 16S rRNA genes were amplified with the V4 amplifying primers 515F and 806R using dual barcoded Illumina primers and Earth Microbiome Project protocols (https://earthmicrobiome.org/protocols-and-standards/16s/ (67)). The resulting 16S rRNA gene amplicons were sequenced at the University of Connecticut Microbial Analysis, Resources, and Services (MARS) facility on the Illumina MiSeq v2.2.0 platform.

Sequence data were demultiplexed and primer and barcode sequences removed by the MARS facility, and then analysed using the mothur package (v. 1.44.1) (68). Briefly, sequences having at least 253 base pairs in length with no ambiguous bases, and no more than eight homopolymer base pairs were retained. Potentially chimeric sequences were identified using the VSEARCH algorithm within mothur (69) using the most abundant sequences in the dataset as a reference, as implemented in mothur, and subsequently removed from further analysis. The remaining sequences were assigned into amplicon sequence variants (ASVs; 100% sequence identity, (70)). Taxonomic classification of AVSs was performed within mothur using the SILVA database (v138 (71)) with the RDP Bayesian classifier (72).

### Statistical and descriptive analyses

All statistical analyses were performed using R studio (v. 2022.12.0(73)). To identify the overall pupation probability, survival curves with 95% confidence intervals were calculated using R packages survival and survminer (74).

16S rRNA gene ASV data was analysed using the phyloseq R software package (75). For the NMDS analyses, Bray-Curtis similarities were calculated for samples and plotted using the plot_ordination command in phyloseq. PERMANOVA statistical testing was performed with the adonis2 function in the vegan R package (76). Alpha diversity was calculated with Shannon’s diversity index and differences in means were determined using an ANOVA test. Post-hoc t-tests were performed with water as a reference level to test if mosquito species differed from water samples. Core ASVs were identified and plotted with the Microbiome R package (77). Differentially abundant ASVs were determined on ASVs found with counts greater than 10 sequences, with unnormalized counts, using the generalized linear model of ALDEx2 and p-vales adjusted for multiple testing using the Bonferroni correction (78) as implemented in the microbiomeMarker R package (79). The ternary diagram was plotted with the ggtern R package (80). The LEfSe analysis (81) to identify potential biomarker strains were also performed within the microbiomeMarker package (79). Differentially abundant taxa between individually and tripartite reared mosquitoes were determined with DESeq2 (82) as implemented in the microbiomeMarker package.

### Data and code availability

The 16S rRNA gene sequence libraries are available in the NCBI SRA under BioProject ID PRJNA943216. Upon acceptance of the manuscript all R code and metadata will be made available within the DRYAD repository.

## ACKNOWEDGEMENTS

This work was funded by a CAES Board of Control Louis A Magnarelli Post-doctoral program grant to BS and DEB. We would like to thank Jacquelyn C. LaReau, John Shepard and Tanya Petruff for technical support.

